# Repeated unilateral injections of botulinum toxin in masticatory muscles in adult rats do not amplify condylar and alveolar bone loss nor modify the volume of the hypertrophic bone proliferation at enthesis

**DOI:** 10.1101/2023.03.10.532076

**Authors:** Pierre Dechaufour, Hélène Libouban, Daniel Chappard, Jean-Daniel Kün-Darbois

## Abstract

**Objectives:** Botulinum toxin (BTX) induces muscle paralysis. It is used in human in masticatory muscles injections performed often repeatedly. A single BTX injection in masticatory muscles in animal induces mandibular bone loss (alveolar and condylar) with a muscle enthesis hypertrophic metaplasia. Our aim was to evaluate mandibular bone changes after unilateral repeated injections of BTX in temporal and masseter muscles in adult rats.

**Materials and Methods:** Mature male rats were randomized into 3 groups: one, two or three injections. Each injection was performed 4 weeks after the prior injection. Each rat received injections in right masseter and temporalis muscles. The left side was the control side. Microcomputed tomography was used to perform 2D and 3D analyses.

**Results:** Bone loss was evidenced on the right sides of alveolar and condylar bone. Alveolar bone volume increased in both control left side and injected right side whereas condylar bone volume remained constant in all groups, for both left and right sides. Enthesis bone hypertrophic metaplasias were evidenced on the BTX injected sides without any modification with the number of injections.

**Conclusions:** BTX repeated injections in masticatory muscles lead to major mandibular condylar and alveolar bone loss that does not worsen. They lead to the occurrence of an enthesis bone proliferation that is not dependent on the number of injections.

**Clinical relevance:** These results are an argument for the safety of BTX injections in masticatory muscles in human.

## Introduction

Botulinum toxin (BTX) is a bacterial metalloprotease produced by an anaerobic bacterium, *Clostridium Botulinum.* There are 7 serotypes of BTX (A to G), but only serotypes A and B are used in clinical practice. The most common serotype is the A. This neurotoxin is a specific ligand of the presynaptic membrane of neurons of the neuromuscular junction. It specifically degrades the SNAP-25 complex of proteins which is necessary for the release of acetylcholine at the axon end. In consequence, acetylcholine is not released from the presynaptic membrane and does not bind to postsynaptic neuromuscular receptors preventing the neurotransmission [1–3]. It then leads to a skeletal muscle paralysis and atrophy which are fully reversible in a few months [4].

The use of BTX in orofacial clinical practice is now well established. It was discovered in the 19^th^ century and was first used, as a therapeutic application, in 1977 in extra-ocular muscles injections for strabismus treatment [5]. BTX is widely used nowadays to treat several diseases with muscular dysfunction such as bruxism, temporo-mandibular joint disorder, orofacial dystonia or myofacial pain [6, 7]. It is also widely used for aesthetic purpose such as the management of facial wrinkles, brow ptosis, excessive gingival display, masseteric hypertrophy, platysmal banding, facial nerve paralysis, hypertrophic scars and keloids [8]. A recent review supports the efficacy and safety of facial BTX injections in clinical practice for bruxism [9]. However, a few studies report mandibular bone consequences, such as a decreased bone density, in human after single or repetitive BTX injections in masticatory muscles [10, 11].

Bone remodeling is a balance between bone resorption and bone formation. Osteocytes are intraosseous cells that are sensitive to mechanical stimuli. They play a major role in the initiation and control of bone remodeling and bone homeostasis [12]. A dysregulation of bone remodeling may occur under the influence of local or general factors. Muscle contractions are necessary for bone homeostasis and a muscle hypofunction leads to bone alterations such as osteopenia [12–16]. It has been shown previously, in animal models, that intramuscular BTX injections lead to long bones osteopenia [14–19]. It was also shown that single BTX injections in masticatory muscles induce mandibular bone loss at alveolar bone and condyle [19–22].

To date only a few studies have investigated condylar articular changes after such BTX injections. Cartilage thickening may occur after BTX injections in growing rats but not in adult rats but this is not constant in the literature which is still controversial on the subject [22–25].

The occurrence of a bone hypertrophic metaplasia that seems to be localized at the enthesis of *Mus. Digastricus* has also been showed to be linked to BTX injections in masticatory muscles of rats [21]. Some cortical bone modifications of the *Mus. Digastricus* enthesis have also been previously reported in human [11]. To our knowledge, this fact has not been investigated by any other author to date and the effects of repeated injections on this bone proliferation have thus not been analyzed. Repeated injections are currently performed in human clinical practice [26]. In consequence, it would be of great interest to investigate the effects of such repeated BTX injections on that animal model.

The aim of the present study was to evaluate mandibular bone and cartilage changes, especially on this bone proliferation, after unilateral repeated injections of BTX in temporal and masseter muscles in adult rats at 1, 2 and 3 months, using microcomputed tomography (microCT) to perform 2D and 3D analyses on condylar and alveolar bone, muscle entheses and condylar articular cartilage.

## Material and Methods

### Animals and experimental procedure

Eighteen weeks-old male Sprague-Dawley rats (n=24), weighing 564 ± 46 g, were used for the study (Janvier-Labs, Le Genest-Saint-Isle, France). They were acclimated for two weeks to the local vivarium conditions (24 °C and 12 h/12 h light dark cycle) where they were given standard laboratory food (UAR, Villemoison-sur-Orge, France) and water *ad libitum.*

Rats were randomized into 3 groups. Rats of group 1 received a single BTX injection (Group 1, n=8), rats of group 2 received two BTX injections (Group 2, n=8) and rats of group 3 received three BTX injections (Group 3, n=8). Each injection was performed with a 4 weeks period after the prior injection. To ensure comparability among each group, all rats were the same age at the first injection. For each rat, the left side was considered as the control side, it did not receive any injection.

BTX injections were performed in each *Mus. Temporalis* and *Mus. Masseter* of the right side only. Each injection procedure was performed as follow: rats were anesthetized with isoflurane and received unilateral injection of type A BTX, 1UI for each muscle (Botox®, Allergan Inc., Irvine, CA, USA); three points of injections for each *Mus. Masseter* and two for each *Mus. Temporalis* were necessary as previously described [21, 22]. Rats were weighed weekly and were sacrificed 4 weeks after their last BTX injection (**Fig. 1**). Facial skin was carefully dissected and removed to perform visual examination of right and left masticatory muscles. Hemimandibles were then dissected, defleshed and fixed in 10% formalin until use.

**Figure 1.**
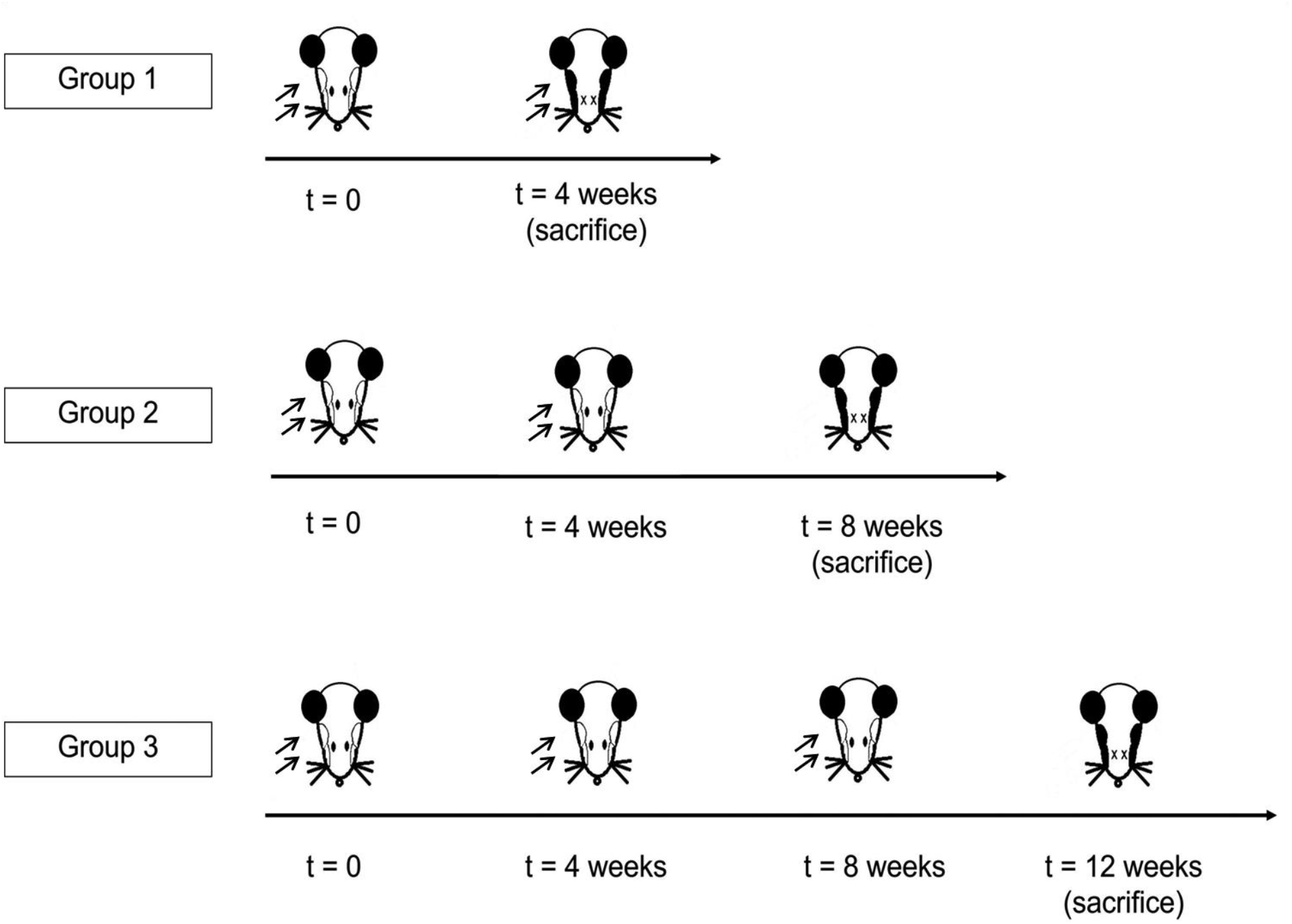
Flowchart of the study

Animal care and experimental protocols were approved by the French Ministry of Research and were done in accordance with the institutional guidelines of the French Ethical Committee (protocol agreement number 01732.01) and under the supervision of authorized investigators.

### Cartilage contrasting procedure

In order to allow articular cartilage analysis, a contrasting method was used as previously described [22]. Briefly, condylar processes were separated from the rest of each hemimandible and immersed for 24 hours in uranyl acetate (UA). UA (Merck KGaA, Darmstadt, Germany) was prepared as a 3% solution in 50° ethanol; the solution was filtered on a 0.2 μm syringe filter and stored at 4°C in the dark. The “stained” specimen was then rinsed during one hour in running tap water and transferred in formalin.

### Microcomputed tomography (microCT)

MicroCT of right and left hemimandibles was performed using a Skyscan 1172 X-ray computerized microtomograph (Bruker microCT, Kontich, Belgium) equipped with an X-ray tube working at 70 kV/ 100 μA. Bones were placed in plastic tubes filled with water to prevent desiccation. The tubes were fixed on a brass stub with plasticine. Analysis was done with a pixel size corresponding to 10.5 μm; the rotation step was fixed at 0.25° with a 0.5 mm aluminum filter. For each hemimandible, a stack of 2D-sections was obtained and reconstructed using Dataviewer software (Bruker) and analyzed using CTan software (Skyscan, release 1.13.11.0).

#### Alveolar and condylar bone measurements

To insure the quality and reproducibility of 3D measurements, reorientation of the alveolar and condylar sections was performed using Dataviewer software. Regions of interest for measurement were then manually selected on several 2D sections to create a 3D volume of interest (VOI) using CTan.

At least ten sections were manually selected for alveolar bone. The anterior limit was the first image where first molar roots and crown were visible. The posterior limit was the last image where third molar crown and roots were observed. Teeth roots, alveolar canal and cortical bone were not included in the VOI which comprised only trabecular bone (**Fig. 2A**).

**Figure 2.**
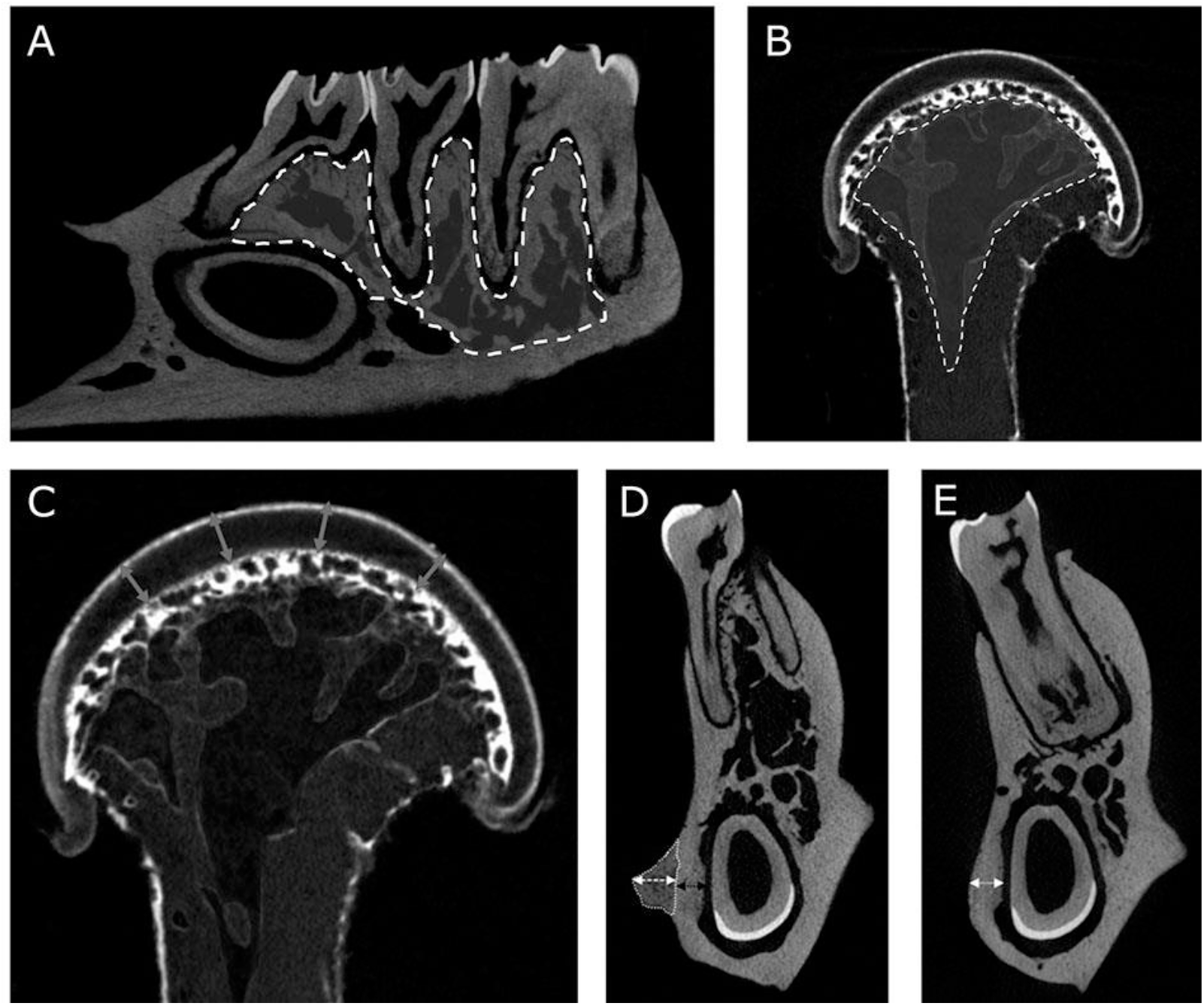
MicroCT analysis. A) View of a region of interest used for 3D alveolar measurement of BV/TV (outlined area by a white dashed line). B) View of a region of interest used for 3D condylar measurement of BV/TV (outlined area by a white dashed line). C) View of a frontal section of right condylar process showing the distribution of the 4 points used for cartilage thickness measurement. D) View of a frontal section of a right hemimandible showing enthesis measurements: enthesis volume (outlined area by a white dotted line), enthesis thickness (white arrow), enthesis cortical thickness (black arrow). E) View of a frontal section of a right hemimandible showing post enthesis cortical thickness measurement.

For condylar bone, height sections were necessary. Anterior limit was the first image of trabecular bone visible when the posterior limit was the last image of trabecular bone visible. Cortical bone and subchondral bone were not included in the VOI (**Fig. 2B**). Trabecular bone volume (BV/TV) was obtained with 3D measurements using CTan. According to guidelines for assessment of bone microarchitecture in rodents using MicroCT, BV/TV is the fraction of the VOI occupied by trabecular bone, expressed in % [27].

#### Condylar cartilage thickness measurements

Condylar cartilage thickness (Cart.Th) measurement was performed on 2D sagittal sections obtained by using the cutting plane facility of the CTan software. For each condyle, 12 measurement points were used, i.e. 4 measurements on 3 different frontal sections. The first frontal section used was the most median section where cartilage thickness seemed the thickest. The other two sections were chosen 125 images ahead and 125 images backwards this first frontal section. These measurements were averaged and expressed in μm for each condyle. Articular cartilage was considered as the area comprised between the tide-mark and the external surface (**Fig. 2C**).

#### Enthesis hypertrophic bone proliferation measurements

The occurrence of a hypertrophic bone proliferation of the *Mus Digastricus* enthesis, on the right side of BTX injected animals, was expected as they had been previously described in this animal model [21]. 2D and 3D measurements of those bone proliferations were performed for each hemimandible using CTan software. The following parameters were measured:

- enthesis volume (in mm^3^), by manually selecting the volume of interest from the first image of visible bone hypertrophy to its last image
- enthesis thickness (in μm), which was defined as the longest height of each enthesis
- enthesis cortical thickness (in μm), which was defined as the thickness of underlying cortical bone.
- post enthesis cortical thickness (in μm), which was defined as the thickness of underlying cortical bone measured at the first distal section without visible bone proliferation (**Fig. 2D-E**).

On left side, where no bone proliferation was expected, two measurements were performed on the corresponding sections using anatomical reference points such as teeth roots, for equivalent enthesis cortical thickness and equivalent post enthesis cortical thickness.

### Statistical analysis

Statistical analysis was performed using the Systat statistical software release 13.0 (Systat Software Inc., San Jose, CA). All data were expressed as mean ± standard deviation (SD). Differences among groups were analyzed by a non-parametric ANOVA (Kruskall-Wallis) and between groups by the Mann and Whitney’s *U* test. Data from right and left hemimandibles were compared using a paired *t*-test. Differences were considered significant when *p < 0.05*.

## Results

### Body weight and anatomic muscle examination

No significant difference in body weight was evidenced within each group during the course of the study except between group 1 and group 3, 4 weeks after the first injections of BTX (rats weighing respectively 559 g and 617g *p*=0.035) (**Fig.3**).

**Figure 3.**
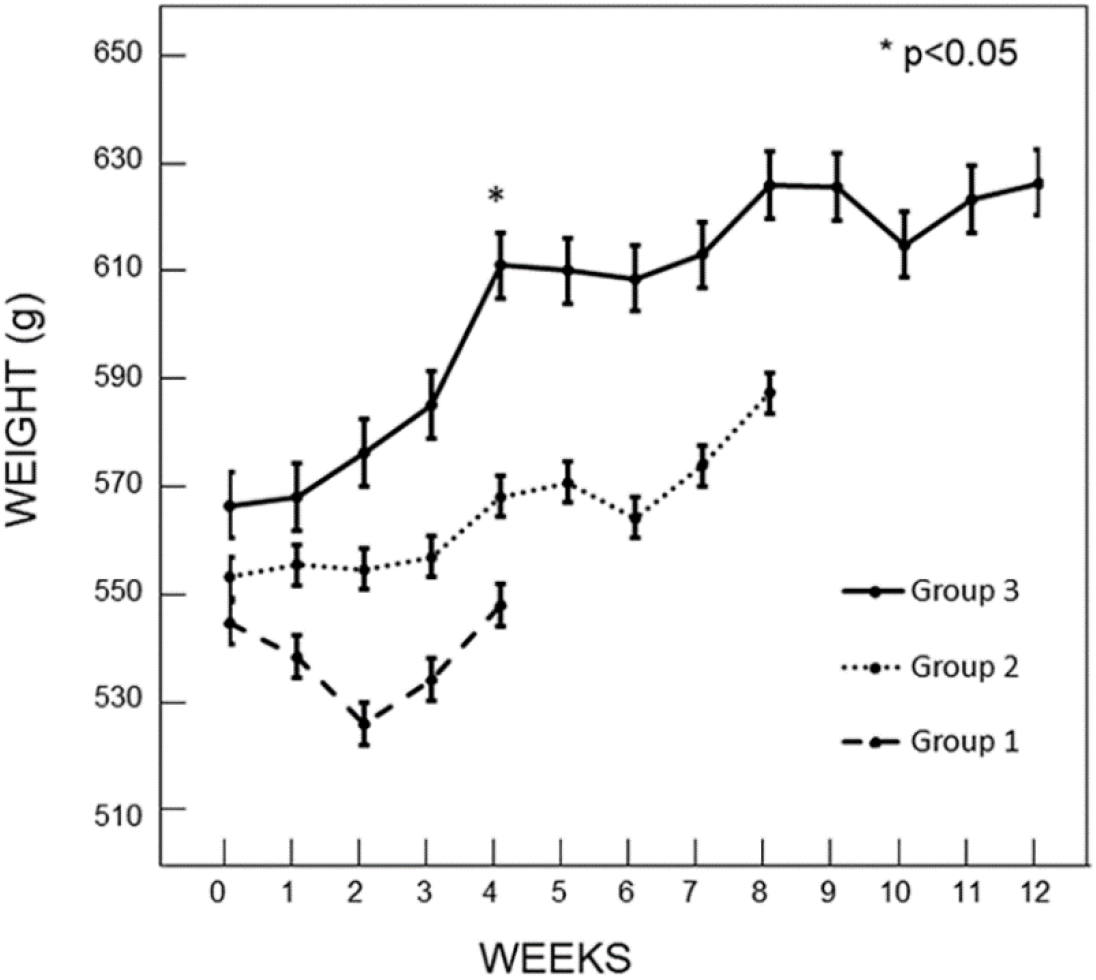
Evolution of body weight showing significant differences between group 1 and group 3 at four weeks. BTX injections were performed at 0 week for all groups, 4 weeks for group 2 and 3 and 8 weeks for group 3 only.

Visual examination during dissection revealed that all rats of all groups presented a *Mus. Masseter* and *Mus. Temporalis* amyotrophy at the right side (**Fig. 4A-B**).

**Figure 4.**
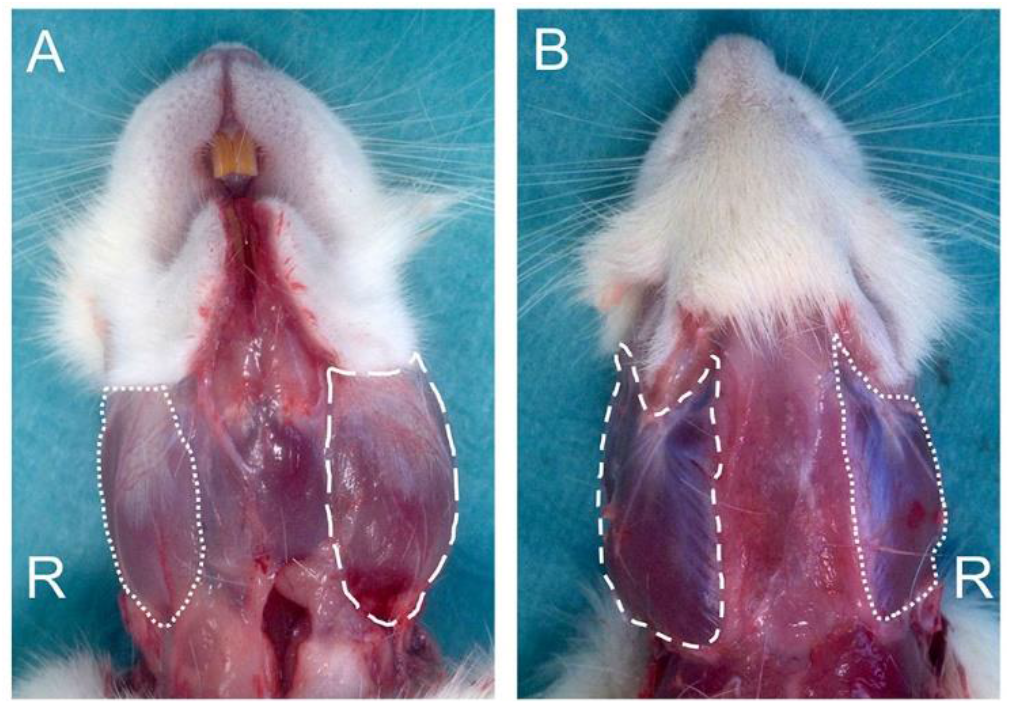
Visual examination and macroscopic aspect of muscles after BTX injections. **A)** in the right *Mus. Masseter* (white dotted line) and left *Mus. Masseter* (white dash line) **B)** in the right *Mus. Temporalis* (white dotted line) and left *Mus. Temporalis* (white dash line).

One rat of group 3 was sacrificed 8 weeks after the beginning of the study. He had started to lose weight 2 weeks before and was sacrificed for ethical purpose since no weight regain had been observed. Post sacrifice examination revealed severely overgrown left superior incisor.

### MicroCT analysis of alveolar and condylar bone measurements

All microCT measurements are summarized in **Table I.** Bone loss was evidenced on the right side of alveolar and condylar bone (**Fig. 5A-B**). Alveolar BV/TV was found reduced in the right side in all group: −12.3%, - 15.3% and −15.1% for group 1, 2 and 3 respectively. Condylar BV/TV was found reduced in the right side in all groups: - 49.8%, - 41.8% and −41.4% for group 1, 2 and 3 respectively. Interestingly a time-related discrepancy could be observed between condylar and alveolar results. No difference within the same sides between group 1, group 2 and/or group 3 could be observed for condylar measurements whereas a significant increase of right and left alveolar BV/TV occurred between group 1 and group 3 (**Fig. 6A-B**).

**Table 1.** Bone morphometric parameters. All data are expressed as mean ± standard deviation (SD).

|  | Group 1<br>left side | Group 1<br>right side | Group 2<br>left side | Group 2<br>right side | Group 3<br>left side | Group 3<br>right side |
| --- | --- | --- | --- | --- | --- | --- |
| BV/TV Alveolar (%) | 52.0 ± 5.0 | 45.6 ± 5.1 <sup>a</sup> | 56.7 ± 4.5 | 48.0 ± 5.3 <sup>a</sup> | 60.7 ± 7.1 <sup>b</sup> | 51.5 ± 6.5 <sup>a, b</sup> |
| BV/TV Condylar (%) | 66.7 ± 6.3 | 33.5 ± 4.5 <sup>a</sup> | 69.4 ± 5.5 | 40.4 ± 3.6 <sup>a</sup> | 67.6 ± 12.9 | 39.6 ± 6.8 <sup>a</sup> |
| Cartilage thickness<br>(µm) | 161 ± 25 | 194 ± 37 | 132 ± 30 | 151 ± 27 <sup>b</sup> | 118 ± 25 <sup>b</sup> | 128 ± 27 <sup>b</sup> |
| Enthesis volume (mm<sup>3</sup>) | 0 | 0.198 ± 0.21 | 0 | 0.172 ± 0.178 | 0 | 0.211 ± 0.131 |
| Enthesis thickness (µm) | 0 | 284 ± 195 | 0 | 240 ± 188 | 0 | 285 ± 111 |
| Enthesis cortical<br>thickness (µm) | 452 ± 52 | 533 ± 95 | 421 ± 55 | 533 ± 95 <sup>a</sup> | 448 ± 35 | 503 ± 57 |
| Post entesis cortical<br>Thickness (µm) | 396 ± 57 | 614 ± 111 <sup>a</sup> | 451 ± 72 | 717 ± 103 <sup>a</sup> | 443 ± 55 | 740 ± 60 <sup>a</sup> |

<sup>a</sup> Indicates a significant difference vs left side of the same group, $p < 0.05$ ; <sup>b</sup> Indicates a significant difference vs group 1 on the same side, $p < 0.05$ .

**Figure 5.**
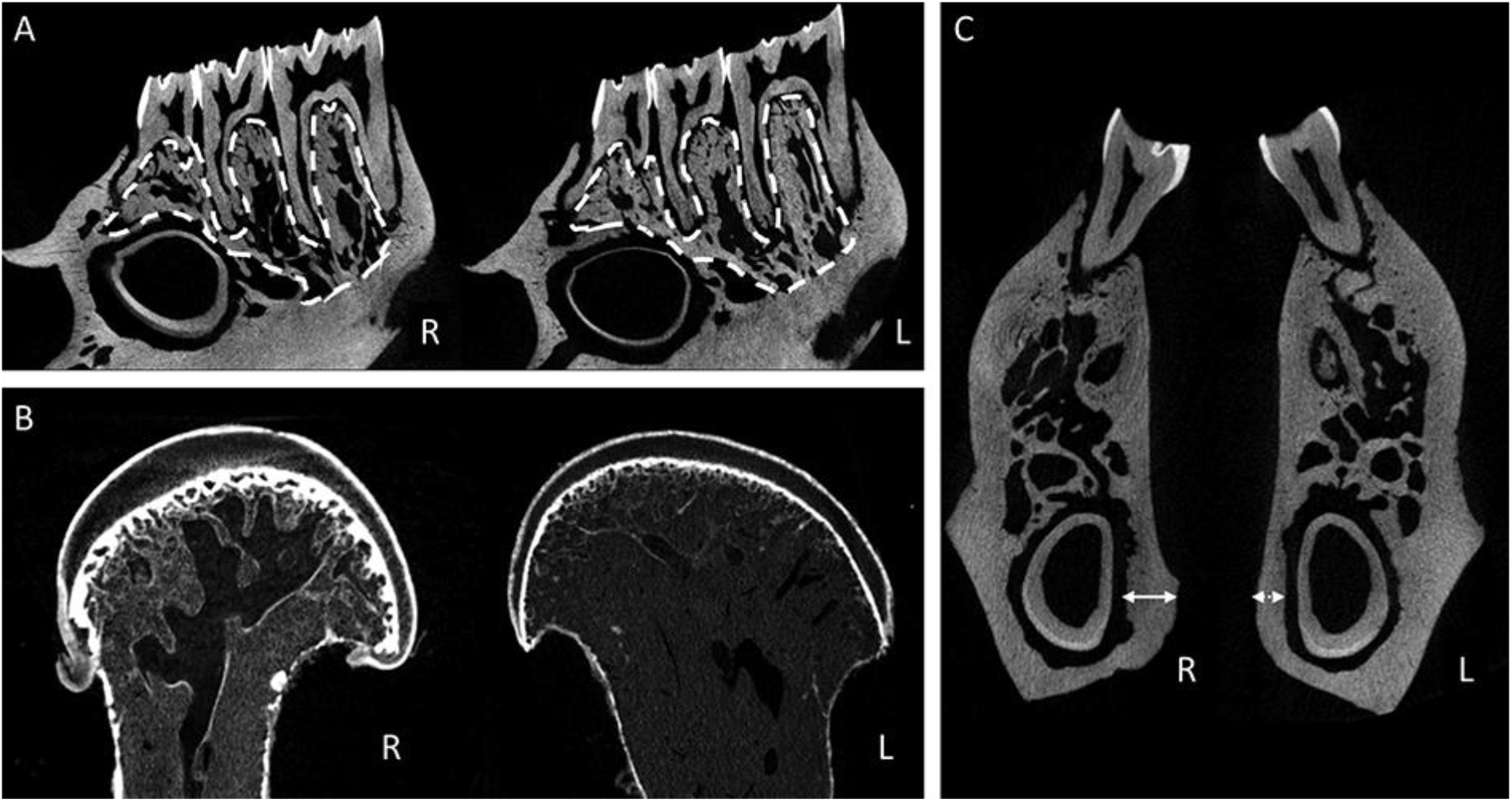
MicroCT analysis as seen on 2D sections. A) Sagittal sections of right (R) and left (L) hemimandibles in a rat from the group 1 showing difference at the alveolar bone; B) Frontal sections of right (R) and left (L) hemi condylar head from a rat from the group 3 showing a marked trabecular bone loss; C) Frontal sections of right (R) and left (L) hemimandibles in a rat from the group 2 at the first distal section without visible enthesis. Note the increasing thickness of the cortical from the right side (white arrow) comparing to left side (white dotted arrow).

**Figure 6.**
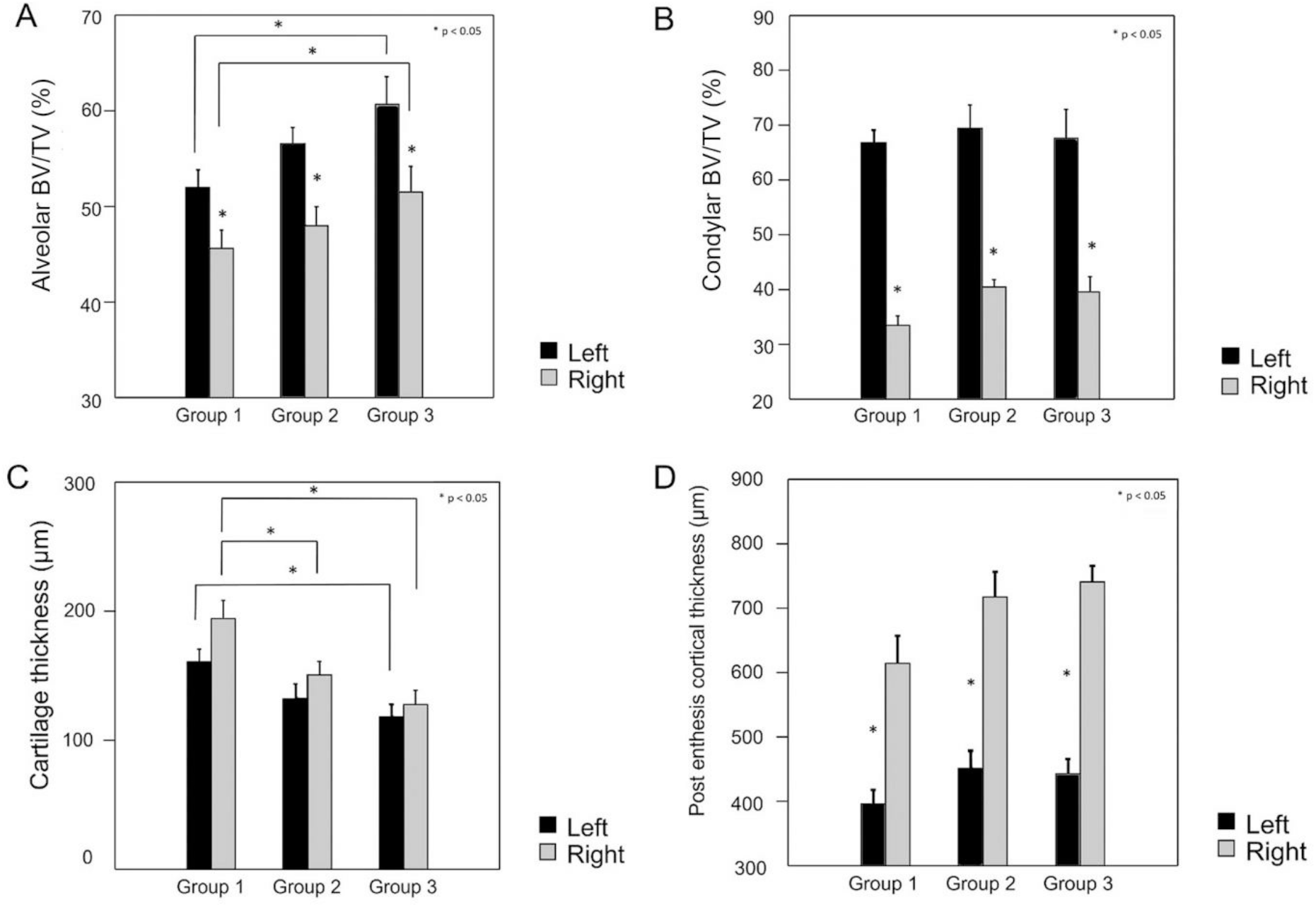
Injected *vs* uninjected side comparisons. **A)** Trabecular bone volume in the alveolar bone (in %) with significant increase of right and left alveolar BV/TV occurred between group 1 and group 3; **B)** Trabecular bone volume in the condylar bone (in %) with a constant reduction at the right sides during the all study; **C)** No significant differences between left side and right side in cartilage thickness, but a significant time related decrease; **D)** Post enthesis cortical thickness was found significantly thicker on BTX injected sides.

### MicroCT analysis of condylar cartilage thickness measurements

After UA impregnation, articular cartilage was clearly visible. No significant difference could be evidenced when comparing cartilage thickness between the right and left sides in each group (**Fig. 6C**). Cartilage thickness significantly decreased with time on both sides reaching a −34% reduction on right side and −27% reduction on left side after three BTX injections.

### MicroCT analysis of enthesis hypertrophic bone proliferation measurements

Hypertrophic bone proliferation at the *Mus. Digastricus* enthesis was only detectable on right side of BTX injected animals. Enthesis volume and thickness remained unchanged with time and number of injections. A significantly thicker enthesis cortical thickness was found on the BTX injected sides only in group 2. However, a trend towards a thicker enthesis cortical thickness was noted in group 1 and group 3, but this difference did not reach significance. Post enthesis cortical thickness was found significantly and largely thicker on BTX injected sides throughout the study (**Fig. 5C and Fig. 6D**). No other difference could be evidenced between and among each group.

## Discussion

Mandibular bone changes after uni or bilateral BTX injections into masticatory muscles in adult and growing animal have been described by a few authors [20]. To date only a few studies have analyzed mandibular bone changes in human after single or repeated BTX injections [11, 28, 29]. The present study aimed at studying the effects of multiple BTX injections on mandibular bone in animal since it had been poorly studied to date [30].

In the present work, the control group was constituted by the non-injected sides (left side). This is usually admitted since previous works on comparable animal models did not show any significant difference between the non-injected and saline injection sides used as control [21, 25, 31].

Rodent incisors grow continuously and must be used regularly to keep them worn down and sharp. If alimentation decreases, then the wear of incisors decreases as well. Insufficient wear of incisors may then result in rapid tooth elongation and thus in malocclusion [32]. Overgrown incisors that are not used enough may thus prevent animals from normal gnawing and lead to undernutrition. In our study, one rat was sacrificed because of an overgrown incisor that led to undernutrition and weight loss. No such case was found in the literature after BTX use. We hypothesize that an imbalance in mastication could explain this fact. However, since only one rat had been concerned we cannot conclude.

Muscular loading plays an essential role in maintaining bone volume and density [33]. Mechanical forces exerted by the masticatory muscles stimulate the mandibular bone and alveolar process to maintain the underlying bone and teeth state. Alveolar bone mineral density is thus considered as correlated with masticatory function [34]. In the present work, a decrease of alveolar trabecular bone volume-on the BTX injected side was found as reported previously in many studies in adult rabbits or rats [21, 31, 35]. Actually, only one comparable previous study does not report such alveolar bone loss in the literature. In this study, only *Mus. Masseter* were injected and animals were sacrificed earlier, probably before any bone tissue changes (i.e., two weeks after injection) [36].

Surprisingly, we evidenced an increase of alveolar bone volume with time and repeated injections of BTX in BTX and control sides. A time-related increase of alveolar BV/TV in standard adult rats cannot be excluded, however it has never been described in the literature. A hypothesis could be that a greater muscle activity occurs in non-injected sides to compensate the lack of muscle activity of BTX side, leading to an increased bone volume with time in non-injected sides. A positive role of normal growth could also be raised to explain this observation, even if rats of the present study were adult rats.

A decrease of condylar trabecular bone volume on the BTX injected side was also evidenced in the present study. Those results are also in line with previous studies [21, 25]. We showed a marked decrease of condylar BV/TV when comparing with alveolar BV/TV. This fact is constantly reported in previous studies with the same animal models [20, 21, 36, 37]. Interestingly, we did not evidence any change with time for condylar BV/TV unlike alveolar BV/TV even if results in adult rats suggest dose dependent deleterious changes in mandibular bone [37]. We can hypothesize this could be due to an earlier bone resorption occurring at the condyles. This seems supported by other studies and could be explained by the presence of secondary cartilage with osteoblasts and osteocytes in the subchondral bone [38]. In addition, studies on rodents have shown a higher mechanical stress in the condylar region [39]. A decrease of almost 30% of BV/TV of the trabecular bone has been evidenced in the condylar region after only one bilateral injections in *Mus. Masseter* of BTX of female rats [40]. Condylar bone loss may have resulted from decreased mechanical loading after BTX injection-induced masseter atrophy.

Articular cartilage thickness decreased after multiple BTX injections. This suggests a time and/or dose dependence link. This is consistent with other previous works where it is evidenced that triple bilateral injections of BTX into the *Mus. Masseter* lead to a greater decrease in cartilage thickness with a reduction in chondrocyte proliferation and differentiation [30]. Mandibular condylar cartilage (MCC) is made of fibrocartilage and it has the unique capacity to adapt to loading changes [41, 42]. After BTX injections, the decrease in *Mus. Masseter* and *Mus. Temporalis* activity may lead to a loading decrease that results in an increased cells apoptosis in the MCC and subchondral bone [43].

An outstanding fact was, once again, the occurrence of a hypertrophic bone proliferation on the BTX injected side of a muscle enthesis as previously described by our group [21]. Enthesis volume and thickness remained unchanged with time and number of injections but it must be noted that enthesis volume and thickness measurements presented high dispersion. The concerned enthesis seems to be the one of *Mus. Digastricus.* It is interesting to note that this muscle is an antagonistic muscle of *Mus. Masseter* and *Mus. Temporalis.* To explain this finding, we hypothesized that a disequilibrium in muscular activity between those muscles lead to an increased bone remodeling at the enthesis of one antagonistic muscle or group of muscles [21, 44, 45]. This bone proliferation occurring in that animal model has not yet been studied by any other authors. The same phenomenon seems to occur also in human after BTX injections [11].

Enthesis morphology seems to depend on the local mechanical environment, i.e., muscle activity, to which it is subjected. It has been showed that functional pressure from another masticatory muscle, the *Mus. Masseter*, greatly affects bone quality as well as the morphological characteristics of its enthesis [46]. In the present work, a heterogeneous and non-time dependent distribution of hypertrophic enthesis volume and thickness and also underlying cortical bone thickness was evidenced. No modification of these parameters occurred with the repetition of BTX injections. A relevant finding was a thickening of the enthesis adjacent cortical bone. This fact cannot be explained and needs to be more investigated. We can still hypothesize that a localized increase muscle activity may lead to hypertrophic proliferation at this muscle’s enthesis and also at the same time to changes in adjacent cortical bone.

## Conclusion

The aim of this study was to investigate mandibular bone changes after repeated injections of BTX in masticatory muscles. A decrease in trabecular bone density of alveolar and condylar regions was evidenced as it had previously been described. However, iterative injections of BTX didn’t amplify bone loss when compared to single injections. Only condylar head cartilage thickness decreased with time and repeated injections. No change in morphologic characteristics of the bone proliferation at the enthesis of *Mus. Digastricus* has been evidenced either. They also seem to be unrelated to the number of BTX injections. An astonishing increase in cortical thickness of adjacent bone has also been found. Those findings, especially the lack of bone loss worsening with time and number of injections, support the use of iterative BTX injections in masticatory muscles in human clinical practice.

## Acknowledgements

Mrs. F. Pascaretti, N. Gaborit and S. Lemière and Mr. J. Roux are thanked for their help.

